# UTILIZING SEQUENCE SIMILARITY NETWORKS FOR CROSS SPECIES ELICITOR IDENTIFICATION OF STREPTOMYCES REGULATORY PROTIENS

**DOI:** 10.64898/2026.05.07.723685

**Authors:** Emilee Patterson, Audrey Birdwell, Andrew Sabatino, Carzon Williams, Allison Walker

**Affiliations:** Department of Biological Sciences, Vanderbilt University, Nashville, TN, USA; The University of Tennessee Health Science Center, Memphis, TN, USA; Hillsboro High School, Nashville, TN, USA; Department of Chemistry, Vanderbilt University, Nashville, TN, USA

## Abstract

*Streptomyces* bacteria produce a variety of secondary metabolites that hold clinical and agricultural value, yet their biosynthetic potential remains unrealized as many biosynthetic gene clusters are not expressed under standard laboratory conditions. Expression of these clusters is tightly regulated, often by cluster situated transcription factors. The TetR family are regulators whose activity is modulated by small molecule elicitors. Although many TetRs have been characterized, elicitors have only been identified for a small fraction of them. This lack of data presents a limitation in our ability to exploit elicitor-regulator pairs for activation of silent clusters and underscores the need for predictive and testable strategies for elicitor identification. In this work, we test the use of sequence similarity networks (SSNs) as a predictor of elicitor identity using the well characterized TetR protein, JadR2, that has a known elicitor, chloramphenicol. We utilized SSNs to identify JadR2 homologs that may also be elicited by chloramphenicol. We developed a heterologous *Escherichia coli* reporter system in which regulator activity was monitored using an EGFP readout of DNA binding activity. Using this system, we screened JadR2 and four homologs for responsiveness to chloramphenicol. We found that 3 homologs were elicited by chloramphenicol, all of which were formerly uncharacterized. These results demonstrate that TetR-family proteins can share elicitor responsiveness and that SSNs can be used to prioritize regulators for functional screening. This work establishes a genomics-informed and bioinformatics-guided framework for linking elicitors to their regulator, expanding the toolkit for natural product discovery by unlocking regulatory information across *Streptomyces*.

## INTRODUCTION

*Streptomyces* are a metabolically prolific genus of bacteria renowned for their ability to produce natural products with a variety of structures and activities(1, 2). These natural products are therapeutically and agriculturally valuable, with activities that treat cancer and microbial infections, or act as pesticides etc.(3–6). However, only a small fraction of *Streptomyces* biosynthetic potential has been realized. Genome sequencing and bioinformatic analyses suggest that a typical *Streptomyces* genome encodes an average of 40 biosynthetic gene clusters (BGCs) with a variety of predicted activities, yet it is estimated that up to 90% of these clusters are not expressed under standard laboratory conditions(7, 8). The mismatch between predicted and expressed BGCs highlights the need for continued research into *Streptomyces* for their biosynthetic potential and continued use as a reservoir for biologically active secondary metabolites.

Advances in genome sequencing and multi-omics approaches have expanded our understanding of *Streptomyces* genomes however, translating this genomic information into chemical output remains a significant challenge(9–11). BGC expression is often tightly controlled by complex regulatory networks that are poorly understood(12). Across all sequenced *Streptomyces* genomes there are hundreds of transcription factors, many of which are uncharacterized. Regulation of secondary metabolism can be highly context dependent, in some cases, the same transcription factor may act differently depending on environmental cues or genomic context, making regulatory outcomes difficult to predict(13–15).

Most BGCs encode one or more cluster situated regulators that govern their expression(16). Traditional strategies to activate silent or cryptic BGCs frequently rely on genetic manipulation of these regulators, such as overexpression of positive regulators or deletion or repression of negative regulators(17, 18). While successful in some cases, these approaches often require extensive cloning and strain engineering that may not be feasible for many *Streptomyces* species. As a result, there is a need for the development of more broadly accessible strategies for targeting and activating secondary metabolism.

One prominent class of cluster situated regulators is the TetR family of transcriptional regulators. TetR proteins are characterized by an N-terminal helix-turn-helix DNA-binding domain and a C-terminal ligand-binding domain(19). TetRs typically bind promoter regions within BGCs and repress transcription. Upon binding to a small molecule ligand, referred to here as elicitors, the protein undergoes a conformational change that reduces DNA-binding affinity, resulting in the activation of transcription(20, 21). Although thousands of TetRs have been described, only a small number of elicitor-regulator pairs have been experimentally characterized(22, 23). The dearth of elicitor data presents a major challenge in exploiting the activity of TetRs for targeted activation of silent BGCs.

Activation of secondary metabolism using elicitors has been explored through untargeted approaches that utilize mass spectrometry to identify changes in the metabolomic profile after treatment with a library of potential small molecule elicitors. The <u>Hi</u>gh-<u>T</u>hroughput <u>E</u>licitor <u>S</u>creening (HiTES) method has been particularly powerful in this context by enabling identification of compounds that induce secondary metabolite production(24). Though robust, these methods do not directly link elicitors to their cognate regulators, making it challenging to generalize findings across species or to predict regulatory activity based on genome sequence alone.

In contrast, directly identifying elicitor-regulator pairs provides a targeted and potentially more generalizable strategy for BGC activation. With the expectation that homologous transcription factors may share regulatory mechanisms, linking specific elicitors to defined regulator families could enable cross-species predictions. If a silent BGC contains a regulator homologous to one with a known elicitor, pathway activation may be possible through treatment with that elicitor, eliminating the need for genetic manipulation or elicitor screening. This approach is particularly exciting for organisms that are difficult to genetically engineer and could offer an alternative option for transcription factor knockouts or overexpression strategies.

Recent work has demonstrated that *Streptomyces* cluster-situated regulators can be functionally reconstituted in heterologous *Escherichia coli* systems as fluorescent biosensors responsive to known signaling molecules(25). These studies establish that pathway associated regulators can retain function outside their native host and highlight the utility of heterologous reporter systems for studying small-molecule sensing. However, these approaches have primarily focused on previously characterized ligand and receptor relationships and have not been widely applied to the discovery of new elicitor-regulator pairs.

In this work, we combine bioinformatics and genomics to predict and validate new elicitor-regulator pairs within the TetR family. Sequence similarity networks (SSNs) are a powerful bioinformatic tool used to group proteins and potentially infer function. In previous work, SSN were used to successfully predict enzyme function (26), here, we expect to use the same tool to predict elicitor-regulator pairs. We constructed a SSN of *Streptomyces* TetR proteins obtained from the antiSMASH database and focused on a cluster containing the well-characterized regulator JadR2, found in the jadomycin B (JdB) biosynthetic gene cluster of *Streptomyces venezuelae*(27). JadR2 represses JdB biosynthesis by binding to the promoter of *jadR1*, a positive regulator of JdB expression. JadR2 has to characterized elicitors, jadomycin B and the antibiotic chloramphenicol (Cm)(28). Binding of either ligand causes JadR2 to release DNA, allowing expression of the JdB cluster.

To test if SSN clusters share elicitor identity, we employed a heterologous *E. coli* expression system in which JadR2 homologs from five *Streptomyces* species were heterologously expressed with an inducible promoter and, on a second plasmid, the *S. venezuelae* JadR2 promotor binding site was cloned upstream of an EGFP reporter. This system enabled direct monitoring of regulator activity and elicitor responsiveness using EGFP expression as a readout of regulator activity while also eliminating the need for native host manipulation. An advantage of this system is that we utilized a known JadR2 operator sequence for functional screening of each homolog. This is both a modular and scalable system that does not require knowledge of the native DNA-binding site for each regulator, resulting in streamlined elicitor identification. By screening JadR2 homologs, we demonstrate that SSNs can successfully predict shared elicitors, providing a framework for targeted activation of silent BGCs through elicitor treatment.

## RESULTS

### Development of a heterologous reporter assay

To assess regulator function, we developed a heterologous reporter system in *E. coli*. This system consisted of a reporter plasmid containing the JadR2 binding site upstream of an EGFP gene and an expression plasmid encoding either JadR2, a homolog, or an empty vector control under IPTG-inducible expression. Double transformants were verified by PCR (Fig S1). EGFP fluorescence normalized to optical density was used as a readout of transcriptional activity (Fig. 1).

**Figure 1.**
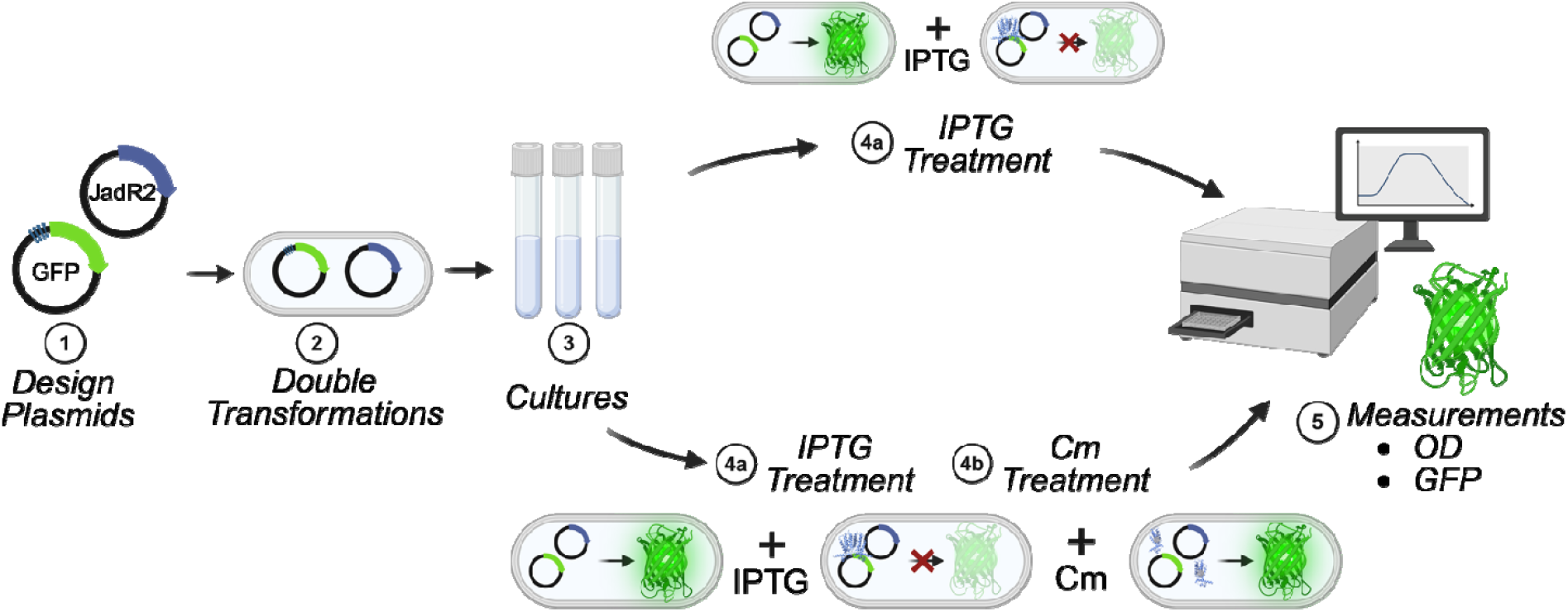
Workflow for identification of elicitor-regulator pairs using a heterologous reporter assay. (1) TetR-family regulators are selected based on sequence similarity to a characterized reference protein and a two-plasmid system is constructed consisting of a reporter plasmid containing a regulator binding site upstream of an EGFP gene and an expression plasmid encoding the regulator under inducible control. (2) Plasmids are double transformed into BL21 (DE3) E. coli. (3) Cultures are grown in EZ Rich Defined Medium overnight. (4a) Cultures treated with IPTG for induction, the expressed regulator binds to its binding sequence upstream of EGFP, repressing expression. (4b) Cultures are treated with IPTG followed by candidate elicitor. Binding of the elicitor to responsive regulators relieves repression, resulting in increased EGFP expression. (5) EGFP fluorescence normalized to OD_600_ is measured as a quantitative readout of regulator DNA-binding activity and elicitor responsiveness.

### Known elicitor-regulator pair, JadR2 and Cm, used as a benchmark for expression and elicitor testing in *E. coli*

Induction of JadR2 expression resulted in a reduction in EGFP fluorescence relative to the empty vector control (Fig 2A). Addition of Cm restored EGFP expression in a concentration-dependent manner (Fig 2B). These results confirm that JadR2 retains both DNA-binding and elicitor-responsive activity in the heterologous system.

**Figure 2.**
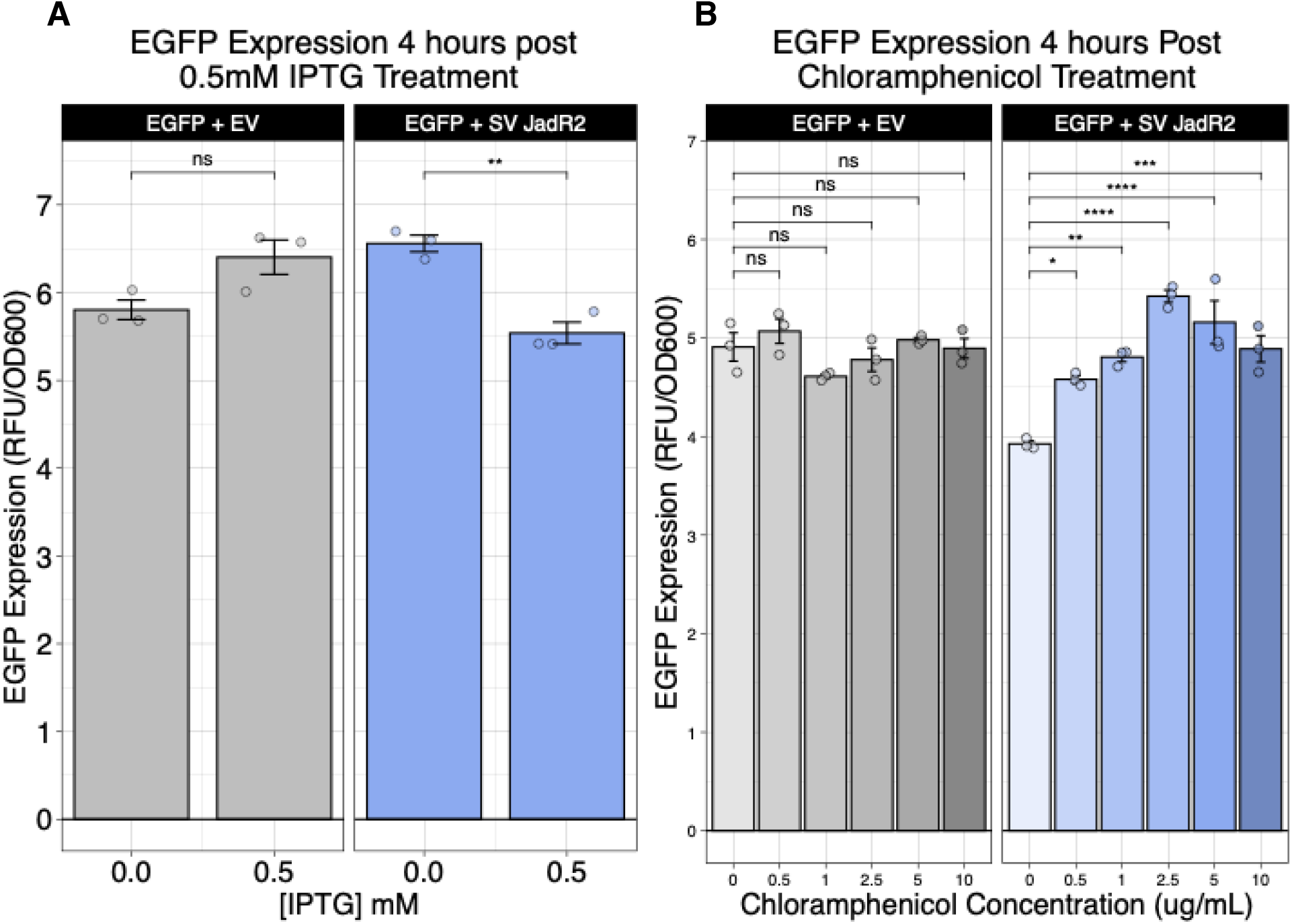
JadR2 retains repression activity and Cm responsiveness in E. coli. The JadR2 protein from Streptomyces venezuelae (SV JadR2) was used as a benchmark for method development. **A**. IPTG-induced JadR2 expression represses EGFP expression. Bar heights reflect EGFP signal normalized to OD_600_ at 4 hours post-IPTG treatment. EV indicates the empty vector control. **B**. Cm treatment restores GFP expression after IPTG induced expression of JadR2. Bar heights reflect EGFP signal normalized to OD_600_ at 6 hours post-CM treatment. EV indicates the empty vector control.

### SSN may be predictive of elicitor identity

To identify candidate TetR-family regulators for elicitor screening, we constructed a SSN using protein sequences from the antiSMASH database (29) and clustered them using the University of Illinois at Urbana-Champaign Enzyme Similarity Tool (30) and Cytoscape (31). Within this network, JadR2 grouped with several protein homologs, suggesting potential functional conservation (Fig 3). Four representative homologs from this cluster were selected for further study based on the percent identity to JadR2. These homologs were derived from *Streptomyces globisporus, Streptomyces exfoliatus, Streptomyces bicolor* and *Streptomyces baarensis*. Sequence alignment revealed conservation within the predicted DNA-binding domain and divergence in regions corresponding to the ligand-binding domain (Fig 4).

**Figure 3.**
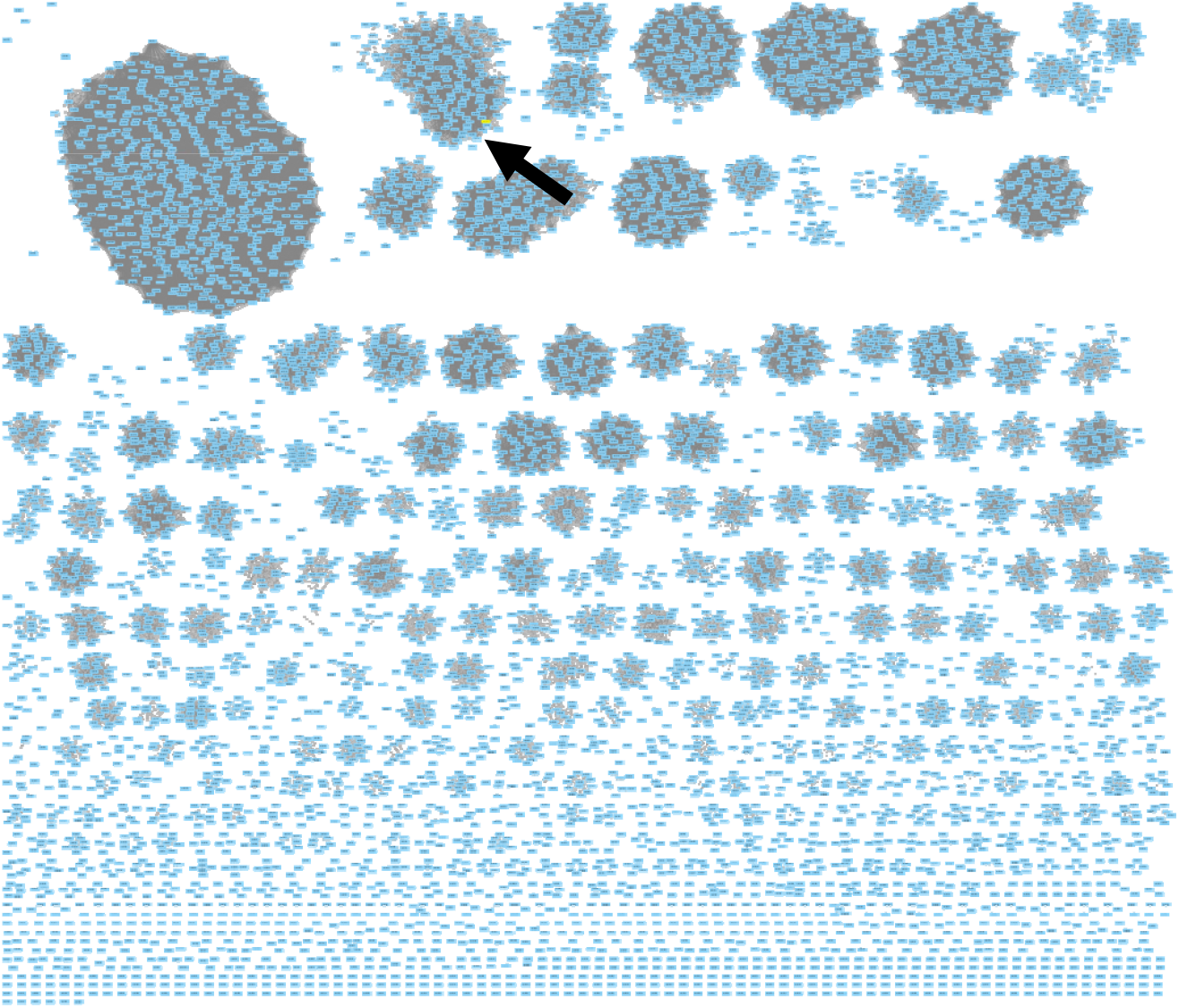
SSN reveals JadR2 homologs selected for use in study. SSN of all TetR proteins from Streptomyces. Arrow pointing to the cluster in which JadR2 from S. venezuelae groups.

**Figure 4.**
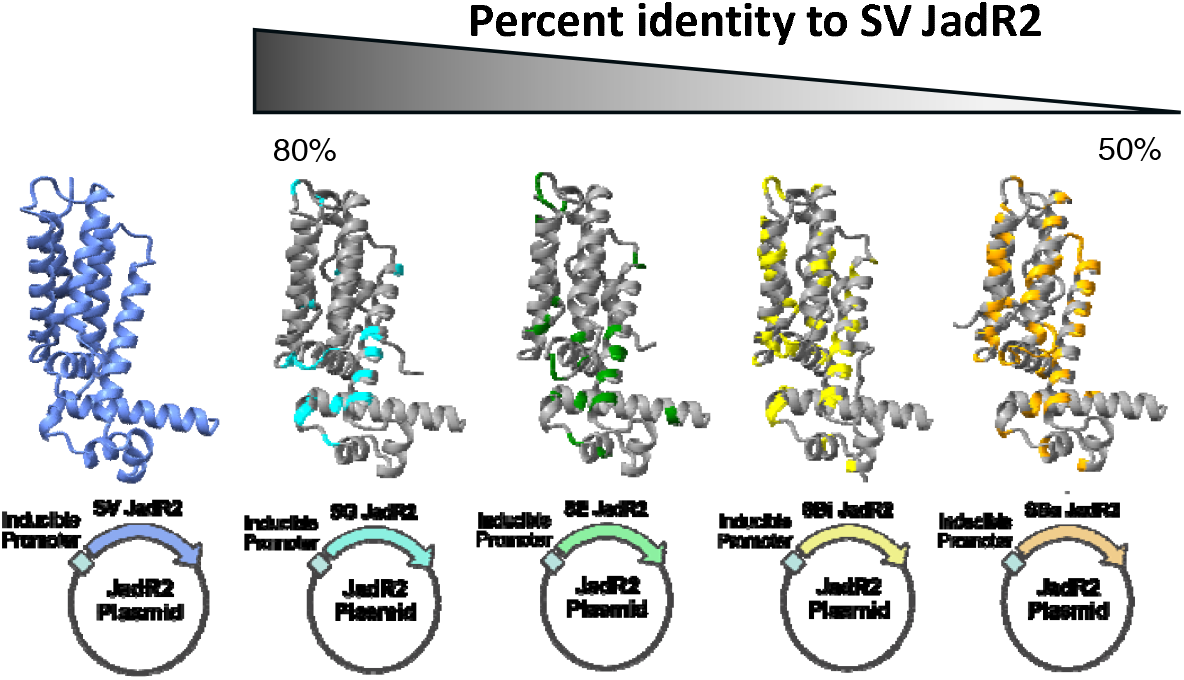
JadR2 Homologs with varying degrees of sequence homology selected. JadR2 homologs from S. globisporus (80% homology), S. exfoliatus (79% homology), S. bicolor (64% homology) and S. baarensis (54% homology) were used for elicitor screening. Protein structure predicted using AlphaFold3 and regions of divergence were mapped in their respective color.

### Homologs of JadR2 selected from SSN for further study

We next evaluated the activity of JadR2 homologs in the reporter system. Four homologs were selected from the SSN JadR2 cluster displaying varying degrees of sequence homolog (Fig 4). Three homologs repressed EGFP expression following IPTG induction, indicating conserved DNA-binding activity (Fig 5A). Treatment with Cm resulted in increased EGFP expression for the same three homologs, although the magnitude of response varied. One homolog did not respond under the tested conditions (Fig 5B). These results identify three previously uncharacterized elicitor-regulator pairs involving Cm responsive TetR homologs. This demonstrates that sequence similarity can be used to prioritize regulators for elicitor screening and supports the use of heterologous reporter systems for functional annotation of transcription factors.

**Figure 5.**
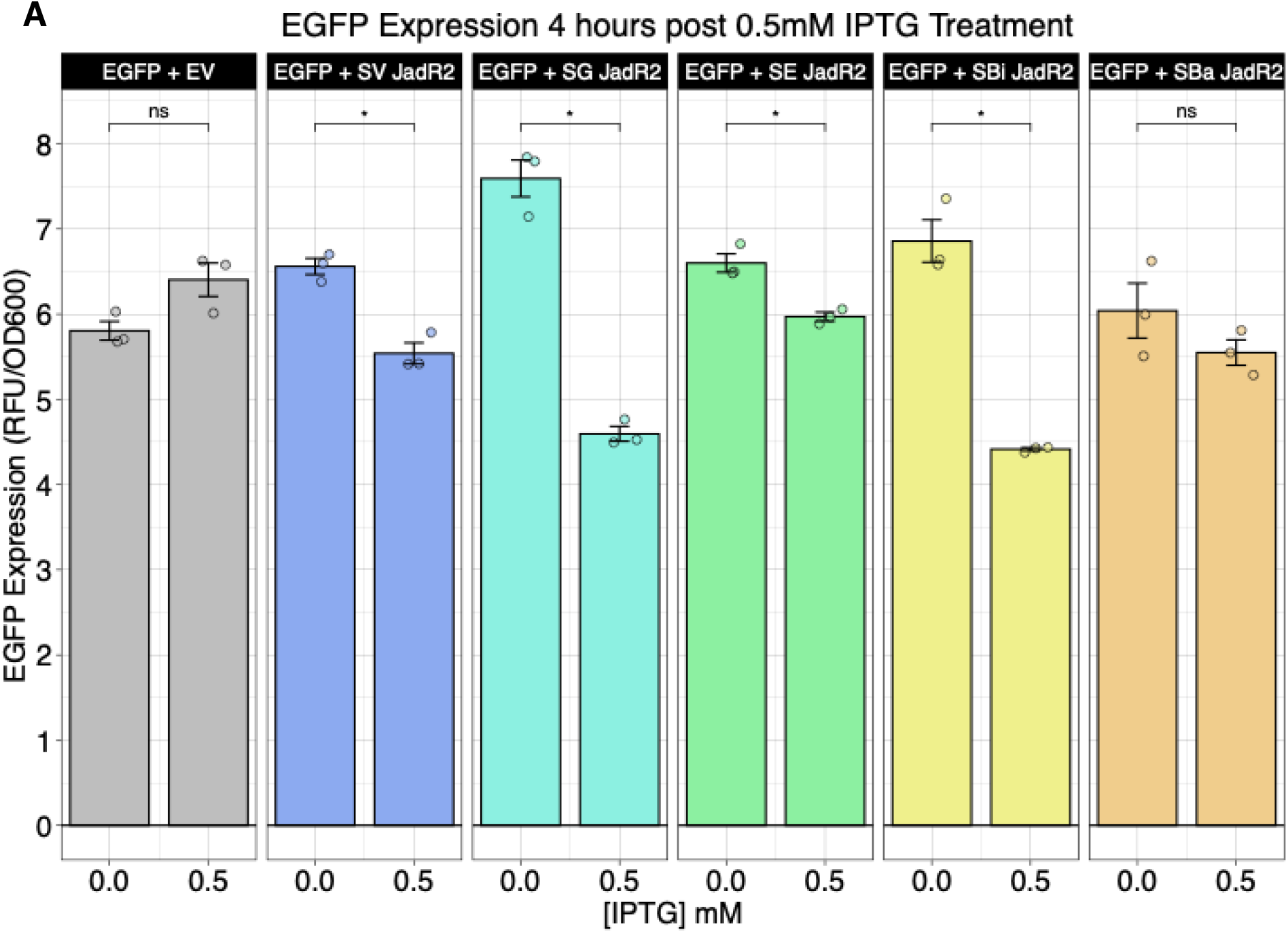
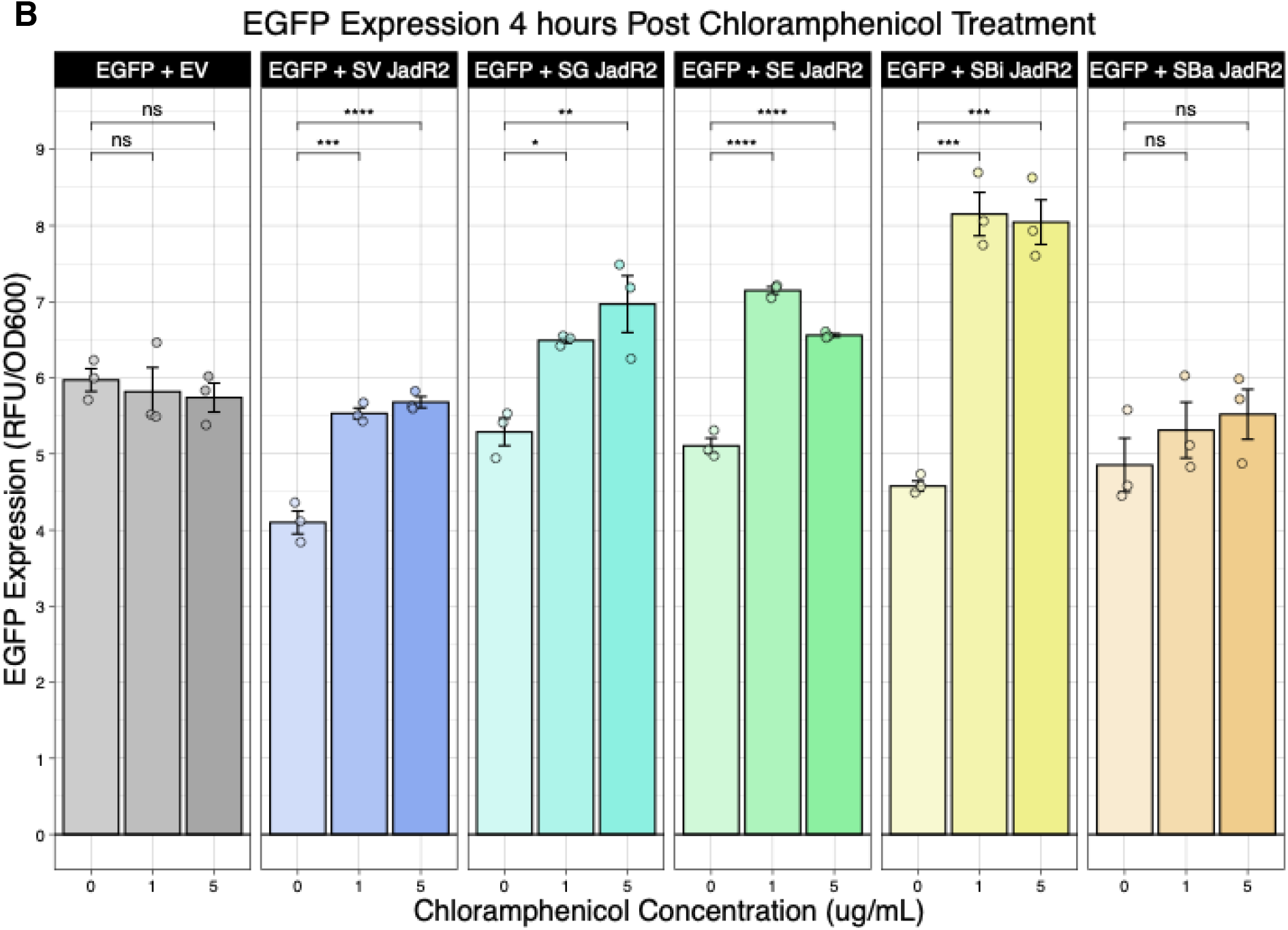
**A**. IPTG-induced JadR2 and JadR2 homolog expression represses EGFP expression via promoter binding. Bar heights reflect EGFP signal normalized to OD600 at 4 hours post-IPTG treatment. **B**. Cm treatment restores EGFP expression after IPTG induced expression of JadR2 and homologs. Bar heights reflect EGFP signal normalized to OD_600_ at 4 hours post-Cm treatment.

## DISCUSSION

In this study, we combined sequence similarity network analysis with a heterologous reporter system to investigate elicitor responsiveness among TetR-family regulators. Using JadR2 as a benchmark, we demonstrate that both repressor activity and elicitor responsiveness are maintained outside the native *Streptomyces* host. Furthermore, three homologs of JadR2 were able to repress transcriptional activity by binding to the native JadR2 operator sequence, characterized by a reduction in EGFP expression. Additionally, those three homologs responded to Cm, characterized by a return of EGFP expression. These experiments expand the number of known elicitor-regulator relationships and highlight the use of SSNs as a predictive tool for elicitor discovery.

These findings support that regulators within biosynthetic gene clusters may be conserved across species, even if the biosynthetic gene cluster they regulate may differ. Hundreds of TetR family regulators are encoded across *Streptomyces* genomes and most lack functional annotation. Strategies that enable prioritization of likely elicitors are essential for leveraging the elicitor-regulator pair relationship to induce expression of silent BGCs, enabling the discovery of novel natural products. Our approach addresses a key limitation of traditional elicitor discovery methods such as untargeted small molecule screening, where changes in metabolite production can be difficult to attribute to specific transcription factors. By directly linking elicitors to their cognate regulators, this work establishes a regulator centered strategy that may allow predictions to extend across species. If homologous regulators share elicitor responsiveness, then small molecules identified in one organism may be used to activate silent biosynthetic gene clusters in related organisms without the need for genetic manipulation or additional elicitor screening campaigns.

This concept is particularly valuable given the challenges associated with genetic engineering in many *Streptomyces* species(32). Traditional approaches to activating silent biosynthetic gene clusters frequently involve overexpression of positive regulators, deletion of repressors, or changing promotors but these methods require cloning, strain engineering, and species-specific optimization that can be time consuming and ineffective. In contrast, elicitor guided activation offers a potentially more accessible alternative in which pathway activation may be achieved through targeted small molecule treatment alone. As genome sequencing continues to reveal large numbers of cryptic biosynthetic gene clusters, approaches that eliminate the need for genetic manipulation could substantially accelerate natural product discovery.

The observation that one JadR2 homolog did not repress EGFP expression or respond to chloramphenicol highlights an important limitation of this strategy. Homologs must be able to bind to the same sequence as a known regulator to reduce EGFP expression to test elicitor specificity. While sequence similarity can narrow the search for regulators with shared function, DNA or ligand specificity may still diverge due to subtle differences in ligand binding domains or regulatory context. This result highlights the importance of experimental validation while also suggesting that sequence similarity networks may be most powerful as prioritization tools rather than definitive predictors of function. Integration of computational transcription factor binding site prediction tools such as COMMBAT(33) may be able to address this limitation by enabling rapid identification of transcription factor binding sites for uncharacterized regulators. Integration of computational DNA-binding site predictions with heterologous elicitor screening could rapid discovery of novel elicitor-regulator pairs from genome sequence alone.

Several limitations of this study provide opportunities for future work. First, this study focused on a single elicitor. Broader chemical screening will be necessary to fully define ligand specificity across homologous regulators. Second, although the *E. coli* reporter system provides a tractable platform for identifying elicitor responsiveness, regulatory behavior in native *Streptomyces* hosts may be influenced by additional transcription factors and other regulatory mechanisms. Third, while the reporter assay measures transcription factor regulatory activity, it does not directly confirm activation of natural product biosynthesis. Future validation in native hosts will therefore be necessary to directly link elicitor treatment to metabolite production.

Future work will expand this framework by incorporating additional regulator families, including GntR, MarR, and LysR regulators, which also play important roles in biosynthetic gene cluster regulation. Expanding SSN guided approaches across regulator families could provide a generalizable strategy for studying elicitor-regulator relationships on a larger scale. In addition, combining this approach with metabolomics and transcriptomics in native hosts will allow direct validation of pathway activation and improve understanding of how elicitor responsiveness translates into chemical output.

More broadly, this work highlights the opportunity to integrate genomics, bioinformatics, and functional screening to better understand microbial gene regulation. As the number of sequenced *Streptomyces* genomes grows, the number of uncharacterized regulatory proteins does as well. Approaches such as the one described here provide a path toward connecting these regulators with the small molecule elicitors that influence them. By enabling transfer of regulatory knowledge across species and reducing reliance on genetic manipulation, this strategy may help access the large reservoir of cryptic natural products encoded in *Streptomyces* genomes.

## MATERIALS AND METHODS

### Sequence Similarity Network Analysis

TetR-family regulator protein sequences were retrieved from antiSMASH annotations from predicted *Streptomyces* BGCs the antiSMASH database (downloaded Aug, 2020). A sequence file was submitted to the Enzyme Function Initiative Enzyme Similarity Tool (EFI-EST), and sequence similarity networks (SSNs) were generated with an alignment score threshold of 40. Networks were visualized and processed in Cytoscape (v3.10.3).

### Plasmid Construction

A reporter plasmid was constructed containing the DNA-binding sequence of interest upstream of EGFP, codon-optimized for expression in *E. coli* (synthesized by Twist Bioscience). Expression plasmids encoding JadR2 or homologs were synthesized (Twist Bioscience) and placed under the control of an IPTG-inducible promoter. Backbone plasmids were pTwist Amp Medium Copy and pET-28a(+), respectively. An expression plasmid lacking an insert was also generated to act as an empty vector control.

The upstream regulatory sequence used in the EGFP plasmid was derived from the intergenic region between *jadR1* and *jadR2* from the jadomycin biosynthetic gene cluster of *S. venezuelae*. To generate SV_Pro_EGFP_E.coli, this *Streptomyces* promoter region was modified upstream of EGFP to optimize expression in an *E. coli* host. The engineered sequence introduced consensus bacterial promoter elements, including a -35 element (TTGACA) and a -10 element (TATAAC) separated by a 17-bp spacer. A Shine-Dalgarno-like ribosome binding site (GAGGAG) was positioned upstream of the EGFP start codon. The EGFP coding sequence was unchanged.

### Bacterial Strains and Transformations

All experiments were performed in *E. coli* BL21 (DE3) strains were grown in EZ Rich Medium (EZRDM), containing 100 μg/ml carbenicillin and 50 μg/ml kanamycin as needed. Reporter and expression plasmids were double transformed using standard protocols. Successful double transformations were confirmed by PCR with plasmid specific primers.

### Fluorescence Reporter Assays

Overnight cultures were grown in EZ Rich Defined Medium (EZRDM) at 37□°C with shaking at 250 rpm. Cultures were diluted 1:10 into fresh EZRDM (5□mL) and grown to mid-log phase (OD_600_ 0.6-0.8). Protein expression was induced with 50□mM IPTG. EGFP fluorescence and OD_600_ were measured in 96-well plate using a plate reader at 2, 3, and 4□h post-induction. Each condition included three biological replicates with and without IPTG. Fluorescence values were normalized to OD_600_ to account for differences in cell density.

### Chemical Elicitor Assays

For elicitor testing, chloramphenicol (0, 1, or 5□mg/mL) was added one hour after IPTG induction. EGFP fluorescence and OD_600_ were monitored as described above.

### Data Analysis and Statistics

Fluorescence data were analyzed after normalization to OD_600._ Biological replicates (n = 3) were averaged, and values are reported as mean ± standard deviation unless otherwise indicated. Statistical analyses were performed in R (v2023.12.1+402). Significance was assessed using two-tailed Student’s t-tests or one-way ANOVA with post hoc correction. Figures were generated using ggplot2 (R package, v3.5.2).

## ACKNOWLEDGEMENTS

This work was funded by a Vanderbilt University Scaling Success award.

