## Supplemental Information for "UTILIZING SEQUENCE SIMILARITY NETWORKS FOR CROSS SPECIES ELICITOR IDENTIFICATION OF STREPTOMYCES REGULATORY PROTIENS"

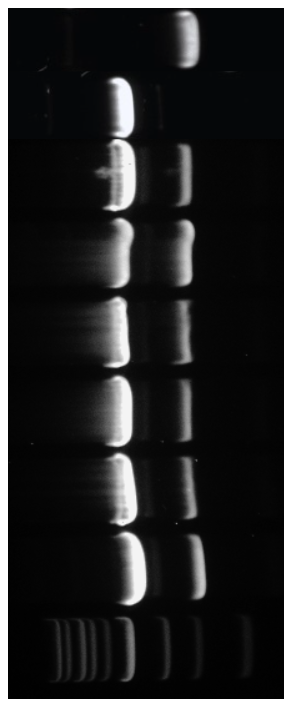

**Figure S1.** PCR verification gel of double transformations containing both the SV_Pro_EGFP_E.coli plasmid and the JadR2 expression vector. Primers specific to each plasmid were used to verify successful transformations. Single transformants of SV_Pro_EGFP_E.coli and SV_JadR2 included as controls to indicate expected band size.

EGFP + SV_JadR2

EGFP +

SG_JadR2

EGFP + SE_JadR2

EGFP + SBi_JadR2

EGFP + SBa_JadR2

EGFP + EV

EGFP

JadR2

| **Plasmid** | **Backbone** | **Insert sequence** |
| --- | --- | --- |
| **SV_Pro_EGFP_E.coli** | pTwist Amp Medium Copy | GAAGTGGTCAAGAGTGCCCGTGGTCTCCCATGGATTCACCTGACTGCTGTTATGTCGCCCCAAGAAACGGACATTCACTGTCAAGTGATGCGATTTCAATCATCACGATGTCAACTCCGTGTCAAATTTTCGTTGCACGACTCTCGGAAAATCTCACACGCCGTGGTAGAGAAATGATTGACACAAACCGGCGCACGGGTTATAACGTTCCCGTTGGCGGGATCCCTACGGGTCCCGGAACTCCGGGACTCTCCAAGGGCCCCGGAAATGAGTGTTAAAGAGGAGAAAGGGAACCATGGTGAGCAAGGGCGAGGAGCTGTTCACCGGGGTGGTGCCCATCCTGGTCGAGCTGGACGGCGACGTAAACGGCCACAAGTTCAGCGTGTCCGGCGAGGGCGAGGGCGATGCCACCTACGGCAAGCTGACCCTGAAGTTCATCTGCACCACCGGCAAGCTGCCCGTGCCCTGGCCCACCCTCGTGACCACCCTGACCTACGGCGTGCAGTGCTTCAGCCGCTACCCCGACCACATGAAGCAGCACGACTTCTTCAAGTCCGCCATGCCCGAAGGCTACGTCCAGGAGCGCACCATCTTCTTCAAGGACGACGGCAACTACAAGACCCGCGCCGAGGTGAAGTTCGAGGGCGACACCCTGGTGAACCGCATCGAGCTGAAGGGCATCGACTTCAAGGAGGACGGCAACATCCTGGGGCACAAGCTGGAGTACAACTACAACAGCCACAACGTCTATATCATGGCCGACAAGCAGAAGAACGGCATCAAGGTGAACTTCAAGATCCGCCACAACATCGAGGACGGCAGCGTGCAGCTCGCCGACCACTACCAGCAGAACACCCCCATCGGCGACGGCCCCGTGCTGCTGCCCGACAACCACTACCTGAGCACCCAGTCCGCCCTGAGCAAAGACCCCAACGAGAAGCGCGATCACATGGTCCTGCTGGAGTTCGTGACCGCCGCCGGGATCACTCTCGGCATGGACGAGCTGTACAAG |
| **SV_JadR2** | pET-28a(+) | GTGACCAAACAAGAGCGGGCCACACGGACGCGCGACGCCCTGATCAAGTCGGCGGCCAGAGAATTCGACGAGCACGGCTACGCCCTGGCCAAGTTGTCGGCGATCAGTTCGGGAGCGGGCGTGAGCCCCGGCGCCCTCCACTTCCACTTCGAGAACAAGGTGGCCGCAGCCGTCGAGATCGACGCGTCCACCACACTGCGCCGGACCGCCAGGATCGTGTACCACCAGCGGTCGAACGCCCTGCAGAACCTCGCCGACACCACCCATGCGCTGGCCCGGCTCGTCCGCGAGGACGTCGTCGTGCGCGCGGGGTTCCGGCTGAGCTGCTCCCAGCTCTGCGGTACGGACCTCAACCTGCGCCAGGAATGGCAGAGCTGCGTCCAGCAGCGGCTCGCGGAGGCCGCCGACGAGGGCCTGCTCGCCTCGGACATCGGTGGTCAACAGGACCTCGCCCGCACGATCGTGGCCGCCACCATCGGCCTGGAGGCCCTCTGCCGGGACAACGGCGAATGGCTGTCGCCCGGCACCGTCACCGGCCTGTGGCGGACCCTCCTCCCCATCGTCGCGGCCCCCGGGCGCTCGCCGCCCTG |
| **SG_JadR2** | pET-28a(+) | GTAACAAAGCAAGAACGTGCGACACGGACGAGAGATGCCCTCATTAGAAGTGCTGCTGGTGAGTTTGATCGACATGGGTATGTTCTTGCTACCTTGAGTGCCATATCAGCCGGCGCCGGGGTCTCCCCAGGGGCTCTGCATTTTCATTTTGAAAATAAAGCGGCTATTGCAGAGGCGGTTGAACAAGAAGCATGTACTGCTCTCCGGCGCACGGCTAGAATTGTTTATGGTCGTCGATCCAATGCGCTTCAAAATCTGACAGATACATCCCATGCATTGGCACGTTTACTGCGAGATGATATTGTGGTACGTGCTGGCTTTCGCCTGAGTTGCGCAGATGATAGCCGCACCGAACTGAGTCTTAGACAAGAATGGCAAAGTTGCGTACAACAACGTCTTGCAGAGGCCGCTGATGAAGGTCTTCTTGCTGCAGGTGTAGGTGGTAGACAAGATCTGGCTCGTACAGTAGTCGCAGCAACCATAGGGTTGGAAGCATTATGTCATGATAATGGTGAGTGGTTATCACCTCGTACGGTTACCGGACTTTGGCGGACACTCCTTCCTATGTTGGCTACACCTGAATCTCTGGCTGGTCTTGACCCTGCTGGTTCGCCTGCAGCAGCAGGAGATGCTCGGGTTCCGGTAGGGTAA |
| **SE_JadR2** | pET-28a(+) | ATGCCTACCAAGGCGCGCCGGATTCCGATTAGTCGTTTCAGAGATGTTGGATGGTATGGAGCCACTGGTAGAAAACCTGTCCCACTTCGTCGTATGGTTTTACGTGAACCATTAGTAACAAAGCAGGAACGTGGGACTCGTACGCGTGATGCACTCATTAGATCCGCCGCAGGTGAATTTGATCTCCATGGTTATGCTCTTGCGACTCTTAGTGCTATTTCATTAGGTGCGGGTGTGTCCCCTGGCGCTCTGCATTTTCATTTTGAAAATAAAGCAGCAATTGCTGAAGCTGTGGAAGATGCCGCATGTACGACACTTAGACGAACTGCGCGTATTGTATATCGGCATAGAAGTAATGCGTTGCAAAATTTAGCAGATACGACTCATGCAATTGCTCGTCTCTTTAGAGAAGATATAGTGGTCAGAGCAGGTTTTCGCTTGTCTTGTGCCAATGATTGTCGGACAGAACTTAATTTGCGTCAAGAGTGGCAATCCTGTGTTCAACAGAGATTAGCTGAAGCAGCTGATGAAGGATTATTAGCTGCAGGGACAGGGCGACAACAAGATTTAGCACGTACCATTGTAGCGGCTACGGTTGGGCTTGTTACTTTGTGTCAAGATTCCGGGGAATGGCTTAGCCCTCATACAGTGACGGGTTTATGGAGAACATTGTTACCAGCGCTGACAGCTCCAGAAGCGTTGGCCGATCTTGAGCCTGCAGGGACTGCTGCAGTTATTACGAGATCCCTGGCTCCTGTGGGTTAA |
| **SBi_JadR2** | pET-28a(+) | ATGACAAAACAAGAGCGTGCGACTAGAACCCGGGATGCCCTTTTGCTTGCTGCTGCGCAAGAATTTCAACGTCATGGCTATGATCGGGCCAAGCTTTCTGCTATTAGTTCGGAAGCTGGCGTATCAAGCGGTGCCTTACATTTTCATTTTGAAAATAAAGCTGCGTTAGCTGAAGCACTCAGAGCGGAAGCTTCTCATGCTCTTCGGCGTGCTGCTCGACTGGCACATCGTCGTAGAACTTCGGCCCTGCAAACCCTTGTGGATTTATCTCATGCTTTAGCCGATGTTTTACGTCGCTCAATTGTGGTGAGAGCAGGTTTTCTTTTGAGTTGTGAAGGTGCAAATGGTACCGATCTCAACTTGCGGCAAGAATGGCAAGTGTGCGTTCAACAACTGTTGGCGGAAGCCGCAGATGAACATCTCTTGGTGCAAGGAGTCCATCAGCAACAACTTACAGGTGTCTTAGTAGCAAGTACAATTGGCTTTGAAGTATTGTCGCGTAGCAATCCAGAATGGTTAAGCTGTTTTACGGTAACAGGATTTTGGGAAGTTTTACTTCCGAATTTGGCTGCGCCGGAAACCTTGACACATCTTGACCCGGCCGGGACCGATACAGCCCATCAAGCTAGCAATGCAGCAGTTAGAACAGGGTTTGTGCCTGCGCCAGCTGGTCCTGCAGCCCCGTAA |
| **SBa_JadR2** | pET-28a(+) | GTTACCAAGCAAGAACGTGCTACACGCACACGGCGTAGTTTGATTCGGTCAGCGGCTGTTGTATTTGAACAACATGGATATGCTCAAGCACGTCTTACCCTTATAAGCTCTGGTGCCGGAGTAAGTACAGGTGCCTTACATTTTCATTTTGAAAATAAAGCAGCAGTGGCCGACGCGGTAGTCGTAGAAGCAAGCCGTGAACTTCGTGAAATGTCAGGGGCTATTAGACGACGTACAGATACTGCATTACAAGCACTGGTTGATTCTTCTCATGCCCTGGCCGAGCGCTTAAGAGAAGATCCTGTTTCCCGTGCAGGTTTTAGACTTTCGTGTGATGCTGCCGGTACTGCAGCACCCGATCTTCGGGTACAATGGCATCATCGGGTTCGTGAGTTATTGGATGATGCCGCGGCGGCAGGGACCTTGGCTGATGATGTGTCACGAGAAGATGCTGCGGCAGCTACAGTAGCTGCTACCGCAGGTTTTGAAGTCCTGGGTCGTCATGACCCGGGCTGGCTCTCTCCATATACCCTTACTGGGTTTTGGCAACTTCTTTTGCCACGTCTGGCAGCAGCCGAAACACTTCCTTTGCTTGATCCCGCAGGAACCGGCGCCACTAGTAGTCCTCGTATACCTGCACCTGCCGCTGGTGCAACGGACGAAGCTGAAGCCGCAGCCGGCGTTGCTAGCGGAGAAGTCGCCAGTGCCGCAGTTGCGGAAGCAGATTAA |

**Table S1.** Summary of plasmid constructs used in this study.

| **PLASMID** | **MEAN**  **(0 IPTG)** | **MEAN**  **(0.5 mM IPTG)** | **p-value** |
| --- | --- | --- | --- |
| *EGFP_SV_JadR2* | 6.5595 | 5.5393 | 0.0035 |
| *EGFP_SG_JadR2* | 7.5913 | 4.5953 | 0.0019 |
| *EGFP_SE_JadR2* | 6.5999 | 5.9698 | 0.0147 |
| *EGFP_SBi_JadR2* | 6.8574 | 4.4141 | 0.0099 |
| *EGFP_SBa_JadR2* | 6.0401 | 5.5481 | 0.2650 |
| *EGFP_EV* | 5.8038 | 6.4038 | 0.0723 |

**Table S2.** Mean EGFP expression values at 0mM and 0.5mM IPTG induction for each plasmid construct with corresponding p values.

*P values were calculated using an unpaired two-tailed Student’s t-test with Benjamini-Hochberg correction for multiple comparisons.*

| **PLASMID** | **COMPARISON** | **p-value** |
| --- | --- | --- |
| *EGFP_SV_JadR2* | 0 to 0.5 Cm | 0.001310901 |
| *EGFP_SV_JadR2* | 0 to 1 Cm | 0.001284004 |
| *EGFP_SV_JadR2* | 0 to 2.5 Cm | 0.00000725525 |
| *EGFP_SV_JadR2* | 0 to 5 Cm | 0.00005425363 |
| *EGFP_SV_JadR2* | 0 to 10 Cm | 0.0005764382 |
| *EGFP_EV* | 0 to 0.5 Cm | 0.8654422 |
| *EGFP_EV* | 0 to 1 Cm | 0.3554508 |
| *EGFP_EV* | 0 to 2.5 Cm | 0.9388478 |
| *EGFP_EV* | 0 to 5 Cm | 0.9950168 |
| *EGFP_EV* | 0 to 10 Cm | 0.9999983 |

**Table S3.** Statistical comparisons of Cm treatment conditions for EGFP+SV JadR2 and EGFP+EV constructs with corresponding p values.

*P values were calculated using a one-way ANOVA with Tukey’s post hoc test.*

| **PLASMID** | **COMPARISON** | **p-value** |
| --- | --- | --- |
| *EGFP_SV_JadR2* | 0 to 1 Cm | 0.0001753 |
| *EGFP_SV_JadR2* | 0 to 5 Cm | 0.0001003 |
| *EGFP_SG_JadR2* | 0 to 1 Cm | 0.0281620 |
| *EGFP_SG_JadR2* | 0 to 5 Cm | 0.0061608 |
| *EGFP_SE_JadR2* | 0 to 1 Cm | 0.0000015 |
| *EGFP_SE_JadR2* | 0 to 5 Cm | 0.0000106 |
| *EGFP_SBi_JadR2* | 0 to 1 Cm | 0.0001053 |
| *EGFP_SBi_JadR2* | 0 to 5 Cm | 0.0001250 |
| *EGFP_SBa_JadR2* | 0 to 1 Cm | 0.6452576 |
| *EGFP_SBa_JadR2* | 0 to 5 Cm | 0.4237825 |
| *EGFP_EV* | 0 to 1 Cm | 0.8877761 |
| *EGFP_EV* | 0 to 5 Cm | 0.9697393 |

**Table S4.** Statistical comparisons of Cm treatment conditions for EGFP+SV JadR2, EGFP+SG JadR2, EGFP+SE JadR2, EGFP+SBi JadR2, EGFP+SBa JadR2 and EGFP+EV constructs with corresponding p values.

*P values were calculated using a one-way ANOVA with Tukey’s post hoc test.*
